# Quantifying and modeling loss of steroid hormones in PDMS-based devices

**DOI:** 10.1101/2025.04.08.647867

**Authors:** Nathaniel G. Hermann, Richard A. Ficek, Dmitry A. Markov, Lisa J. McCawley, M. Shane Hutson

## Abstract

Many polydimethylsiloxane (PDMS)-based devices, *e*.*g*., organ-on-chip or microphysiological systems, have been developed to investigate biological processes at a miniaturized scale. These devices typically culture cells under microfluidic perfusion to dynamically dose cells with chemicals of interest; however, PDMS is known to interact with hydrophobic compounds and can strongly limit such compounds’ in-device bioavailability. Here, we quantify chemical-PDMS interactions for three commonly used steroid hormones: aldosterone, estradiol, and progesterone. We find that aldosterone does not detectably interact with PDMS; estradiol interacts modestly; and progesterone interacts strongly. Based on these measured interactions, we computationally model dynamic dosing protocols based on pulsed/bolus delivery and circadian control. We show that interactions with PDMS can strongly disrupt these dynamic dosing protocols in a chemical-specific and flow-rate-dependent manner. These results have critical implications for the use of steroid hormones in PDMS-based devices.

## Introduction

Many miniaturized biomedical devices, including lab-on-a-chip, organ-on-a-chip (OOC), and microelectromechanical systems (MEMS) are fabricated from polydimethylsiloxane (PDMS); however, PDMS is known to interact with hydrophobic chemicals. When using such devices to study the dose-response curve of a drug or toxicant, researchers must mitigate chemical-PDMS interactions and/or account for those interactions to predict actual in-device doses [1]. In a similar vein, when culturing cells in PDMS-based devices, users need to know whether hydrophobic components of the cell culture media could be significantly depleted by partitioning into PDMS. For example, early studies showed that cells cultured in PDMS-based devices were insensitive to added estrogen because it was sequestered by PDMS and no longer bioavailable [2]. Here, we investigate and quantify chemical-PDMS interactions for three key steroid hormones – aldosterone, estradiol, and progesterone – which are all commonly used as growth factors in cell culture media [3, 4, 5, 6, 7]. These hormones are also a direct or indirect target of study for OOC/MEMS-based research in reproductive biology and medicine, *e*.*g*., in endometrium-on-a-chip or mammary-gland-on-a-chip devices [8, 9, 10, 11, 12, 13, 14, 15, 16, 17, 18, 19, 20].

The need to quantify and model hormone-PDMS interactions is further exacerbated by several use cases with dynamic hormone dosing. In the simplest cases, a pulse or step change in hormone levels may be used to trigger a change in cell behavior or differentiation [21, 22, 23, 24, 25], but reversible interactions with PDMS could yield reduced-amplitude pulses with long tails of low-level dosing. In more complicated scenarios, an experiment may require the modulation of hormone levels to match circadian rhythms [26] or menstrual cycles [8, 27]. In these periodic cases, interactions with PDMS can alter both the amplitude and phase of hormone dosing. Finally, for an experiment that measures the cellular secretion of a hormone over time, an appropriate model of hormone-PDMS interactions is needed to convert measured concentrations to dynamic secretion rates. There are mitigation strategies, such as using cell culture media with fetal bovine serum (FBS), that can stabilize the concentration of hydrophobic growth factors [28], but these strategies also alter dosing dynamics. They are also not amenable to applications that require serum-free media.

The three steroid hormones tested here are structurally similar, but their properties differ enough to expect them to interact with PDMS to different degrees. Aldosterone is a mineralocorticoid, estradiol is an estrogen, and progesterone is a progestogen, but their structures all share a gonane core. They also all share similar molecular weights (360.4, 272.4, and 314.5 amu) [29] and collisional cross sections (181.09, 173.34, and 182.2 Å^2^) [30]. The key difference is that aldosterone is much less hydrophobic than estradiol or progesterone, as demonstrated by their logP values of 1.08, 4.01, and 3.87 [29]. Additionally, both aldosterone and estradiol have two hydrogen bond donors, while progesterone has none. Having higher logP and fewer hydrogen bond donors have both been implicated as key properties that increase a chemical’s propensity to interact with PDMS [31]. As shown below, and as expected based on these characteristics, aldosterone interacts weakly, if at all, with PDMS while estradiol interacts moderately and progesterone interacts very strongly.

## Methods

### PDMS Preparation

PDMS Sylgard 184 (Dow Corning, Auburn, MI) was mixed in a 10:1 mass ratio of elastomer base to curing agent. PDMS membranes were spun out from small volumes of PDMS to 80-µm thickness on a Laurell WS-400-6NPP Spin Coater (Laurell Technologies Corporation, Lansdale, PA) using the procedure described by Markov et al. [32], and then cured overnight in a 67°-C oven.

### Hormone Preparation

Aldosterone, estradiol, and progesterone were acquired as powders from Sigma Aldrich (St. Louis, MO). Stock solutions were prepared in neat dimethyl sulfoxide (DMSO) at approximately 70 mM concentration. Aqueous solutions were then prepared via a dilution of DMSO stock in 1x pH-7.4 phosphate buffered saline (PBS) (Thermo Fisher, Waltham, MA) to concentrations between 5 and 0.1 mM.

### Diffusion-through-Membrane Experiments

To determine chemical-PDMS interactions, we follow the procedure for diffusion-through-membrane experiments as described in Hermann et al [1]. In brief, two chambers separated by an 80-µm thick PDMS membrane were each filled with 200 µL of solution – one containing a hormone solution (source) and one containing a blank solvent mixture (sink). Each chamber was then sampled multiple times over 24 hours and UV-Vis spectroscopy was used to track hormone concentrations in both source and sink. Interaction parameters were estimated by fitting the dynamic source and sink concentrations to a mass-transport model, with numerical solutions to the model generated in COMSOL Multiphysics (COMSOL Inc., Burlington, MA) and fitting conducted in Mathematica (Wolfram Inc., Champaign, IL).

### Simulations of Dynamic Hormone Exposures

To estimate how PDMS interactions would modulate dynamic hormone exposures, we used COMSOL Multiphysics to perform computational fluid dynamics and reaction-diffusion simulations using a finite element method [1]. Simulations were run for aqueous solutions of both progesterone and estradiol flowing through PDMS microchannels under pulse-chase and circadian rhythm scenarios.

## Results

As shown in Fig. 1A-B, both progesterone and estradiol partition into and diffuse across an 80-µm thick PDMS membrane, with the source and sink chambers approaching equilibrium over hours to days. In contrast, aldosterone does not cross the membrane (Fig. 1C). To quantitatively characterize chemical-PDMS interactions for progesterone and estradiol, we fit this data to a partition-diffusion model [1] to determine four key parameters: the effective diffusivity in solution (*D*_*S,eff*_); diffusivity in PDMS (*D*_*P*_); a mass-transfer coefficient at the solution-PDMS interface (*H*); and the PDMS-solution partition coefficient (*K*_*PS*_). To get spectroscopically detectable concentrations of estradiol and progesterone into solution, we conducted these experiments with 5-30% DMSO as a cosolvent. We used a previously derived log-linear relationship [1] to extrapolate from the partition coefficients in mixed solutions, *K*_*P S*_, to estimate the partition coefficients in pure aqueous solution, *K*_*P W*_. That relationship is

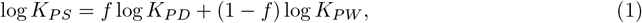

where *f* is the volume fraction of DMSO and *K*_*P D*_ is the PDMS-DMSO partition coefficient. The log-linear fits to the data are shown in Fig. 1D. The best-fit values of *K*_*P W*_, *K*_*P D*_, and *D*_*P*_, plus a non-rate-limiting lower bound for *H*, are shown in Table 1. We find that progesterone partitions into PDMS two orders of magnitude more strongly than estradiol, and diffuses within PDMS about one order of magnitude faster.

**Figure 1.**
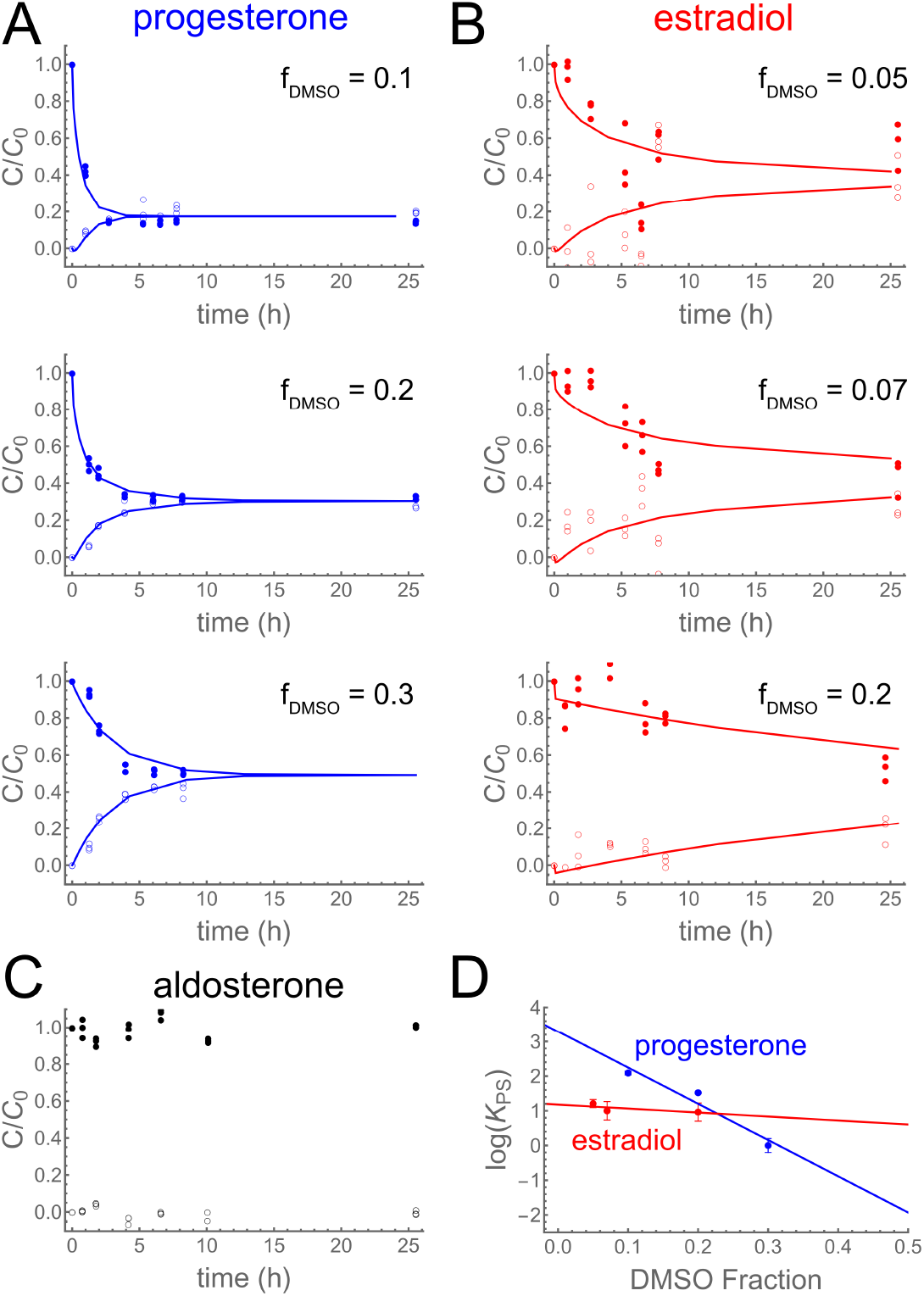
Hydrophobic steroid hormones partition into and diffuse through PMDS. (A-C) Diffusion-through-membrane results for (A-blue) progesterone, (B-red) estradiol, and (C-black) aldosterone. Filled symbols are concentrations in the source chamber; open symbols are concentrations in the sink chamber; lines are best fits to a partition-diffusion model. For progesterone and estradiol, the DMSO co-solvent fraction is as shown (increasing top to bottom). (D) Extrapolation of progesterone and estradiol results to zero DMSO based on a linear regression of log *K*_*P S*_ versus DMSO fraction.

**Table 1.** Chemical-PDMS interaction parameters for steroid hormones.

| Chemical | $\log K_{PW}$ | $\log K_{PD}$ | $\log D_P$ (mm<sup>2</sup>/h) | $\log D_S$ (mm<sup>2</sup>/h) | $\log H$ (mm/h) |
| --- | --- | --- | --- | --- | --- |
| progesterone | $3.30 \pm 0.60$ | $-7.20 \pm 0.60$ | $-1.90 \pm 0.40$ | 0.336 | $\geq 1.56$ |
| estradiol | $1.18 \pm 0.14$ | $+0.03 \pm 0.14$ | $-3.03 \pm 0.32$ | 0.306 | $\geq 1.56$ |
| aldosterone | — | — | — | 0.296 | — |

Note that these experiments were conducted on a blot mixer to keep solutions well mixed. The fits thus estimate effective diffusion coefficients in solution, *D*_*S,eff*_, which are not generally applicable for further device simulations. We instead report calculated values of *D*_*S*_ based on the methods of Miyamono and Shimono [33] with van der Waals radii estimated using the molecular descriptor package Mordred [34, 35]. These values of *D*_*S*_ (1.98 - 2.17 mm^2^/h) are about an order of magnitude smaller than the self-diffusivity of water (10.8 mm^2^/h) [36] and are reported as log *D*_*S*_ values in Table 1.

### Simulations

To investigate and illustrate the impact of steroid-PDMS interactions, we used finite-element modeling to simulate the distributions of progesterone and estradiol in two dynamic, microfluidic use cases. We first simulated 1-hour pulses of each hormone as delivered through a rectangular microchannel: 10.55-mm long by 1.5-mm wide by 100-µm tall and embedded in a 4-mm thick slab of PDMS. These simulations were performed with volumetric flow rates, *Q*, of 1, 10 and 100 µL/min. As shown in Fig. 2 (top), the slowest of these flow rates yielded the strongest departures from the input square pulse. For strongly-interacting progesterone at that flow rate, less than 0.1% of the chemical reaches the end of the microchannel. For moderately-interacting estradiol, almost all the chemical reaches the end of the microchannel, but its interactions with PDMS round off the square pulse and lead to an extended delivery of hormone for about half an hour after the nominal end of the pulse. At faster flow rates, the delivered pulses of estradiol are only slightly rounded off, but those for progesterone remain limited to about 40% (*Q* = 10 µL/min) or about 80% (*Q* = 100 µL/min) of the input dose. Progesterone also has a low-dose, post-pulse delivery that lasts for a few hours.

**Figure 2.**
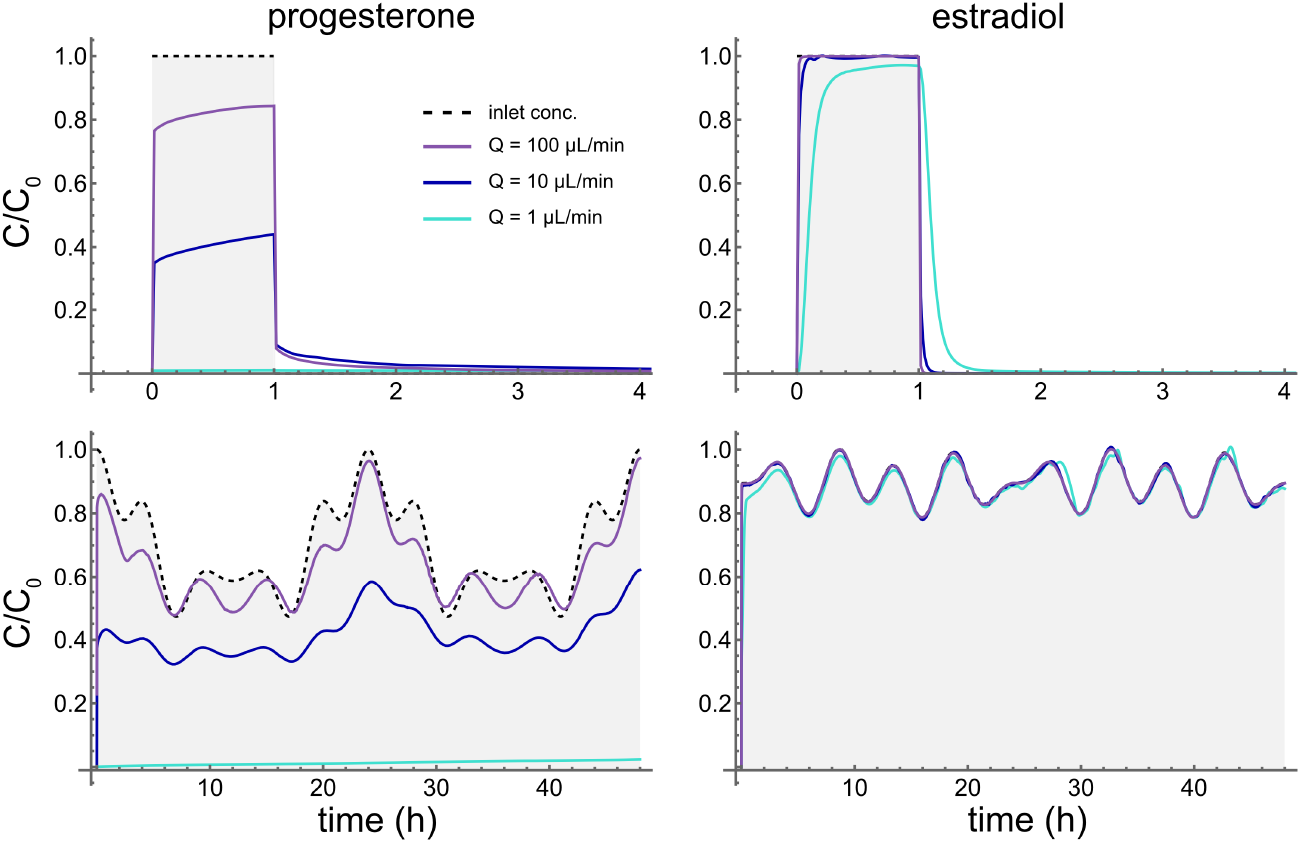
The fidelity of dynamic dosing protocols is disrupted by chemical-PDMS interactions in a flow-rate-dependent manner: (left column) progesterone; (right column) estradiol; (top row) simulations of a 1-hour pulsed dose; and (bottom row) simulations of circadian variations in hormone levels. Shaded regions with dashed outlines correspond to inlet concentrations. Colored lines correspond to concentrations at the channel midpoint for volumetric flow rates of 1, 10 and 100 µL/min (colors as shown in the legend).]

For the second dynamic use case, we simulated delivery through the same microchannel but with an input concentration that follows a prescribed circadian rhythm. The different rhythms chosen for progesterone and estradiol are Fourier-smoothed versions of data from an *in vivo* study by Bugnam et al [37]. As shown in Fig. 2 (bottom), moderately-interacting estradiol can be delivered similarly to the targeted circadian rhythm at all tested flow rates. In contrast, strongly-interacting progesterone can be delivered somewhat closely to the targeted rhythm only at the highest flow rate (100 µL/min). Decreasing the flow rate to 10 µL/min reduces the circadian variation by about half; decreasing to 1 µL/min completely eliminates the ability to follow the prescribed dynamics.

## Discussion

As we have shown, both estradiol and progesterone interact with PDMS, while aldosterone does not. Given that aldosterone has a logP of 1.08, this result is consistent with previous reports that chemicals with logP *≤* 1.8 do not partition strongly into PDMS [31]. The other two steroids have logP well above this threshold and do interact with PDMS, albeit in distinct ways. Progesterone strongly partitions into PDMS (*K*_*P W*_ *∼* 1990) and diffuses through bulk PDMS at a moderate rate (*D*_*P*_ *∼* 0.013 mm^2^/h). Estradiol partitions more weakly (*K*_*PW*_ *∼* 15.2) and diffuses through PDMS more slowly (*D*_*P*_ *∼* 9.33 *×* 10^*−*4^ mm^2^/h).

Simulations of pulsed or bolus delivery of these steroid hormones reveal the key role of flow rate in determining the concentration lost in a channel. At low flow rates (*Q* = 1 µL/min), progesterone is strongly depleted from solution (only 0.03% is bioavailable at the end of the 10.55-mm channel), and the pulse of estradiol is rounded off and extended by a slow release of hormone from the microchannel walls well after the end of the pulse. Higher flow rates shift the system into advection-dominated transport, resulting in much more progesterone and nearly all estradiol reaching the target. Using higher flow rates can thus mitigate the effects of chemical-PDMS interactions, but only if cells in the device can tolerate the higher flow rates.

Simulations of circadian hormone delivery with progesterone and estradiol suggest that in-device dosing requires careful management to account for chemical-PDMS interactions. At low flow rates (*Q* = 1 µL/min), interactions with PDMS severely limit in-device dosing of progesterone (less than 2.4% of targeted exposure after 48 hours) and dampen any periodic behavior. At faster flow rates (*Q* = 10 *−* 100 µL/min), more than 30% of the progesterone is delivered through the channel, but its periodic behavior is phase-shifted. Further, in the case of 10 µL/min flow rates, the amount of progesterone delivered increases with each 24-hour cycle. On the other hand, weaker interactions with PDMS allow allow nearly complete delivery of estradiol in a circadian rhythm at all tested flow rates. These simulations raise concerns about the feasibility of circadian (or menstrual) control of hormone concentrations in PDMS-based devices with naive aqueous solution or serum-free media. For strongly-interacting chemicals and physiologically-relevant flow rates, it seems likely that the addition of serum, carriers, or co-solvents will be necessary to avoid severe hormone depletion or phase-shifting.

Taking a broader view of the state of research in PDMS-based devices, we note that the steroids discussed here are a tiny subset of the cell culture components that could interact strongly with PDMS. Typical culture media include a range of hydrophobic hormones, vitamins, lipids, and growth factors.

These are often introduced as components of added serum, and while the carrier proteins in serum do help mitigate loss of these hydrophobic compounds into PDMS [28], the composition of the serum is often poorly defined. For PDMS-based devices, it may be advisable to use chemically-defined, serum-free media. With a well-defined medium and measured chemical-PDMS interaction parameters, one can use 3D chemical transport modeling to ensure that the in-device concentrations of relevant compounds are known and controlled.

## Conclusion

We have determined that the steroid hormones estradiol and progesterone both partition into and diffuse through PDMS; a third less hydrophobic steroid, aldosterone, does not. By parameterizing the interactions of estradiol and progesterone with PDMS, we have developed simulations of pulsed or bolus delivery and circadian concentration control in PDMS-based devices: Both dosing schemes can be compromised by hormone-PDMS interactions. Researchers interested in applying dynamic dosing of these hormones in PDMS-based microfluidic systems can use finite-element modeling to account for and plan strategies to mitigate in-device hormone loss.

## Data availability

Data for this article, including the results of membrane experiments and simulated solutions used in generating interpolation functions, are available at Open Science Framework at https://osf.io/z5bvx/.

## Acknowledgements

This publication was supported by U.S. Environmental Protection Agency (EPA) STAR Center Grant #84003101. Its contents are solely the responsibility of the grantee and do not necessarily represent the official views of the U.S. EPA. Further, U.S. EPA does not endorse the purchase of any commercial products or services mentioned in the publication.

